# Xanomeline restores endogenous nicotinic acetylcholine receptor signaling in mouse prefrontal cortex

**DOI:** 10.1101/2022.09.29.510152

**Authors:** Saige K. Power, Sridevi Venkatesan, Evelyn K. Lambe

## Abstract

Cholinergic synapses in prefrontal cortex are vital for attention, but this modulatory system undergoes substantial pre- and post-synaptic alterations during adulthood. To examine the integrated impact of these changes, we optophysiologically probe cholinergic synapses *ex vivo*, revealing a clear decline in neurotransmission in middle adulthood. Pharmacological dissection of synaptic components reveals a selective reduction in postsynaptic nicotinic receptor currents. Other components of cholinergic synapses appear stable, by contrast, including acetylcholine autoinhibition, metabolism, and excitation of postsynaptic muscarinic receptors. Pursuing strategies to strengthen cholinergic neurotransmission, we find that positive allosteric modulation of nicotinic receptors with NS9283 is effective in young adults but wanes with age. To boost nicotinic receptor availability, we harness the second messenger pathways of the preserved excitatory muscarinic receptors with xanomeline. This muscarinic agonist and cognitive-enhancer restores nicotinic signaling in older mice significantly, in a muscarinic- and PKC-dependent manner. The rescued nicotinic component regains youthful sensitivity to allosteric enhancement: treatment with xanomeline and NS9283 restores cholinergic synapses in older mice to the strength, speed, and receptor mechanism of young adults. Our results reveal a new and efficient strategy to rescue age-related nicotinic signaling deficits, demonstrating a novel pathway for xanomeline to restore cognitively-essential endogenous cholinergic neurotransmission.

## Introduction

Even in healthy aging, there are declines in sustained attention [1,2], cue detection [3,4], and executive function [5,6]. These abilities all rely on cholinergic control of the prefrontal cortex [7], with basal forebrain projections releasing acetylcholine in deep cortical layers [8–11]. Not surprisingly, it is hypothesized that age-related cognitive changes arise from disruption of the cholinergic system [12–14]. Human clinical investigations reveal age-related reductions in basal forebrain volume [15,16], decreases in nicotinic binding [17,18], and possible changes in muscarinic binding [19,20] and acetylcholinesterase [21,22]. Targeting the cholinergic system has shown some promise in treating age-related cognitive changes in humans [23–26][27–29][30–32]. However, treatment strategy is limited by the substantial knowledge gap in the integrated functional impact of age-related changes to prefrontal cholinergic synapses. This uncertainty complicates the identification of specific treatments capable of restoring youthful functional parameters.

Similar to humans, mice show progressive cognitive changes that occur during adulthood [33,34], beginning as early as 4-5 months [35]. Transgenic mice have been used to investigate the complexity of cortical cholinergic synapses [36–39], revealing the functional roles of multiple pre- and post-synaptic receptors and regulatory enzymes [36–39]. However, previous work has not examined the impact of age on these complex synapses. The need to study the integrated impact of adult age-related changes on endogenous cholinergic signaling is underscored by research suggesting that cholinergic receptor expression is dynamic [40,41] and there may be interactions across receptor subtypes [42–44].

Here, we use optogenetic tools to investigate the impact of age on integrated functioning of prefrontal cholinergic synapses from young to middle adulthood. We demonstrate a striking and selective reduction of the postsynaptic nicotinic optophysiological response while revealing the stability of pre- and postsynaptic muscarinic responses. We discover that the age-related nicotinic functional deficit is not amenable to positive allosteric modulation. Therefore, we harness signaling pathways of the well-preserved muscarinic receptors as an unorthodox approach to nicotinic potentiation. Overall, we demonstrate the strength, speed, and receptor mechanism of cholinergic signaling can be restored. Xanomeline provides a new and efficient strategy to rescue age-related nicotinic signaling deficits.

## Materials and Methods

Methods are summarized briefly below. See **Supplemental Methods** for additional description.

### Animals

The University of Toronto Faculty of Medicine Animal Care Committee approved all experiments in accordance with the guidelines of the Canadian Council on Animal Care (protocol #20011621). We performed experiments on transgenic mice expressing channelrhodopsin in cholinergic afferents (ChAT-ChR2; RRID:IMSR_JAX:014546, Jackson Laboratory [45]). This transgenic line labels cholinergic neurons and avoids labeling neurons only transiently cholinergic in development [46]. This approach also avoids use of anesthetic agents known to perturb nicotinic receptors [47,48] and surgery known to impact cognitive function [49]. A total of 60 mice were used for this study (age range: ~P50-350), with balanced numbers of males and females. This age range was selected because mice show progressive, age-dependent changes in behavior through adulthood [33–35].

### Electrophysiology and Optogenetics

For whole-cell patch-clamp electrophysiology, Layer 6 pyramidal neurons in the prelimbic and anterior cingulate regions of the medial prefrontal cortex were identified by size, morphology, and proximity to white matter. Responses from a homogenous population of layer 6 regular spiking pyramidal neurons [39] were examined in voltage-clamp at −75mV and in current-clamp.

To excite chanelrhodopsin-containing cholinergic afferents, blue light (470nm) was delivered in 5 ms pulses with an LED (2mW; Thorlabs) through at 60x objective lens. As developed previously in our lab, this opto-ACh stimulus is delivered in a frequency-accommodating (50Hz to 10Hz) eight pulse train to stimulate cholinergic axons [39] to mimic the activation pattern of cholinergic neurons [50–52].

### Experimental design, analysis, and statistics

Mice of both sexes were used across ages and in pharmacological interventions. Experiments draw from multiple mice per intervention, and recordings were made from 1-3 neurons per brain slice, with 2-3 slices per mouse. Unless otherwise stated, pharmacological agents were pre-applied for 10 minutes and co-applied during optogenetic stimuli and recovery period. Magnitude of cholinergic currents were measured from exponential fit by peak amplitude (pA) and charge transferred (pC) into the cell, measured by area under the current response for 5s from opto-ACh onset. Opto-ACh responses plotted in a histogram by age were analyzed statistically by nonlinear robust regression (Graphpad Prism 9.3.1). Age differences in the presence of pharmacological acetylcholine blockers were analyzed with unpaired or paired, two-tailed *t*-tests, where applicable. Pharmacological manipulations to improve nicotinic responses were evaluated with respect to age by two-way ANOVA and one-way ANOVA, where applicable.

## Results

### Postsynaptic responses to optogenetic release of endogenous acetylcholine decrease with age

To stimulate release of endogenous acetylcholine (opto-ACh, **Fig 1A,B**), we use a pattern of optogenetic stimulation [39] designed to mimic burst firing with spike frequency accommodation seen in cholinergic neurons innervating the prefrontal cortex [50–52]. With whole cell electrophysiology, we measure opto-ACh responses in regular spiking pyramidal neurons in layer 6 with voltage-clamp at −75 mV. The brief opto-ACh protocol elicits inward current responses resilient to combined application of glutamatergic synaptic blockers CNQX (20μM) and D-APV (50μM), (peak amplitude, opto-ACh: 54.5 ±21.22pA, opto-ACh+synaptic blockers: 55.5 ±18.5pA; paired *t*-test, *t_6_* = 0.2, *P* = 0.9) and sensitive to combined cholinergic receptor antagonists atropine (200nM) and DHβE (3μM), (0 ±0% opto-ACh current remaining with cholinergic blockers; paired *t*-test, *t_21_* = 4.8, *P* = 0.0001). This pharmacological assessment suggests opto-ACh currents are directly mediated by cholinergic receptors on the recorded pyramidal neurons.

**Figure 1.**
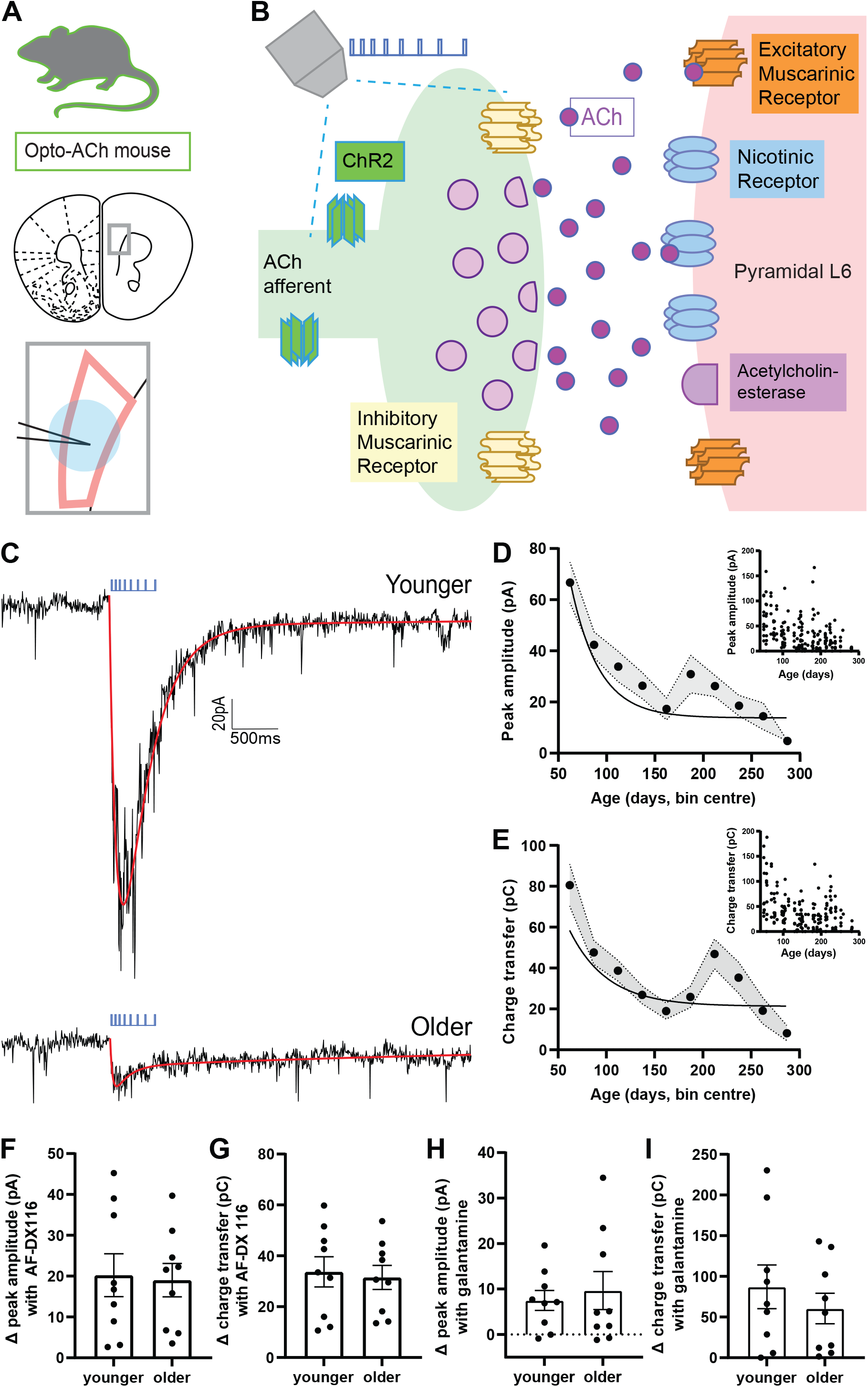
Responses to optogenetically-released acetylcholine decrease with age. (**A**) Image depicts coronal brain section taken for whole-cell electrophysiological recordings. The area outlined in pink shows layer 6 of the prefrontal (prelimbic and infralimbic) cortex. The recording electrode is represented in black and the delivered optogenetic stimulus is represented in blue. See caveats section for further discussion of the model of opto-ACh stimulation. (**B**) Schematic represents the cholinergic prefrontal synapse receiving optogenetic stimulus (0.5s pulse train of decreasing frequency). Afferents release endogenous acetylcholine (ACh) which binds to nicotinic and muscarinic receptors, exciting postsynaptic pyramidal neurons. (**C**) Example recordings of opto-ACh current from a neuron from younger and older mice, measured in voltage clamp (Vm = −75mV, stimuli in blue, triple exponential fit in red). (**D, E**) Graphs show the decrease with age of the peak amplitude and the charge transfer of the opto-Ach current (n = 185 neurons, 43 mice). (**F, G**) Graphs show the increase in opto-ACh current peak and charge transfer with M2 muscarinic antagonist AF-DX 116 are not different between younger (< P150) and older (≥ P150) responses (younger: P87± 16.1, P50-130; older: P225 ± 32.2, P159-303). (**H, I**) Graphs show the change in peak amplitude and charge transfer of the opto-ACh current with galantamine are not different between younger and older responses (younger: P104 ± 18.7, P57-138; older: P202 ± 16.1, P150-281).

To explore age-related changes in this functional cholinergic signal, we record whole-cell responses to opto-ACh across a broad age distribution in adulthood. The vast majority of regular spiking pyramidal neurons (91.3%) respond to opto-ACh with an inward current ≤ 5 pA that can be fit to an exponential curve for quantification (**Figure 1D, E)**, with all neurons included in the graphs. We find a significant decline in opto-ACh response with age (**Fig 1C,D,E)**, as measured by peak amplitude of the response (R^2^ = 0.165; *P* = 0.0001, **Fig 1D** *inset*) and the area-integrated measure of charge transfer (R^2^ = 0.141; *P* = 0.0004, **Fig 1E** *inset*). Fitting a nonlinear robust regression to peak amplitude and charge transfer reveals a fast decline in opto-ACh responses in early adulthood before responses plateau into middle adulthood and beyond (**Fig 1D, E**).

Since 5 months of age in mouse is associated with the emergence of cognitive changes [33–35] and an inflection point in our analysis of response amplitude and charge transfer appears at about P150 (**Fig 1D,E**), we chose to contrast responses from above and below this age. This comparison reveals that the “older” middle adulthood mice (P150 and older) had an opto-ACh response ~50% lower than those of “younger” adult mice in terms of peak amplitude (younger, 42.7 ±3.5pA; older, 22.6 ±2.8pA, unpaired *t*-test, *t_183_* = 4.5, *P* < 0.0001) and charge transfer (younger, 48.6 ±4.1pC; older, 27.8 ±2.8pC, unpaired *t*-test, *t_183_* = 4.2, *P* < 0.0001). From this grouped perspective, we find the decline in cholinergic neurotransmission occurs in the absence of commensurate changes in postsynaptic neuron intrinsic properties (**Supplemental Table S1**), although the older group showed a modest increase in resting membrane potential (~2mV; unpaired *t*-test: *t_213_* = 3, *P* = 0.004). Since our dataset included both males and females, we also considered the effect of sex on the opto-ACh responses (**Supplemental Figure S1**). There was no significant interaction between sex and age group by two-way ANOVA. There was an effect of sex, with greater responses seen in females (F_1,180_ = 4.6, *P* = 0.03). The difference is consistent with existing work showing sex differences in layer 6 pyramidal cholinergic responses [41,53]. However, it is a modest difference compared to the strong and significant effect of age group, where opto-ACh responses are decreased in older relative to younger mice (F_1,180_ = 23.5, *P* < 0.0001). Since sex does not appear to alter the age-related cholinergic change, both sexes are included in further experiments to examine cellular mechanisms underlying decreased opto-ACh response with age.

To probe components of the cholinergic synapse that may contribute to the age-related decline, we interrogate the functional role of the autoinhibitory muscarinic M2 receptor in the opto-ACh response. Located presynaptically, inhibitory M2 receptors modulate cholinergic responses [54–56], restricting responses to opto-ACh at the prefrontal synapse [39]. We find that blocking muscarinic-mediated autoinhibition with muscarinic M2 receptor antagonist AF-DX 116 (300nM) significantly increased opto-ACh responses peak amplitude (paired *t*-test, *t_17_* = 4.7, *P* = 0.0004) and charge transfer (paired *t*-test, *t_17_* = 8, *P* < 0.0001) in neurons across the age range. However, there were no significant age-related differences in the increase in peak amplitude (unpaired *t*-test, *t_16_* = 0.2, *P* = 0.9; **Fig 1F**) nor charge transfer (*t_16_* = 0.3, *P* = 0.8; **Fig 1G**). For examples demonstrating the impact of blocking M2-mediated autoinhibition on younger and older opto-ACh responses, see **Supplemental Fig S2A**. These results suggest that muscarinic autoinhibition exerts a consistent effect on synaptic cholinergic signaling across the age range examined.

To further examine synaptic components that exert control over cholinergic responses, we consider the functional role of acetylcholinesterase in the opto-ACh response. Investigations of acetylcholinesterase in pyramidal neurons of post-mortem human tissue show changing levels in aging, without consensus on direction of change [22,57] or examination of the functional impact of these changes. We show that blocking acetylcholinesterase with inhibitor galantamine (1μM) significantly amplified and prolonged opto-ACh response peak amplitude (paired *t*-test, *t_16_* = 3.7, *P* = 0.002) and charge transfer (paired *t*-test, *t_16_* = 4.6, *P* = 0.0003) in neurons across the age ranged examine. However, there were no significant age-related differences in increase in peak amplitude (unpaired *t*-test, *t*_15_ = 0.5, *P* = 0.7; **Fig 1H**) nor charge transfer (unpaired *t*-test, *t*_15_ = 0.8, *P* = 0.4; **Fig 1I**). For examples demonstrating the impact of inhibiting acetylcholinesterase on younger and older opto-ACh responses, see **Supplemental Fig S2B**. These results suggest that acetylcholinesterase activity exerts a consistent effect on synaptic cholinergic signaling across the age range examined.

Both presynaptic autoinhibition and acetylcholinesterase suppression at prefrontal cholinergic synapses are intact through adulthood, indicating that the observed decline in opto-ACh responses may have a postsynaptic origin.

### Postsynaptic selectivity: age-related decline in nicotinic signal, stable muscarinic signal

To identify whether postsynaptic nicotinic and muscarinic components of opto-ACh responses each show an age-dependent decline, we pharmacologically isolated these two optophysiological signals in separate experiments. Our strategy was based on evidence that the opto-ACh response is abolished by co-application of atropine and DHβE, as shown above and in [39]. Therefore, we examined the isolated nicotinic component after muscarinic blockade with atropine and the isolated muscarinic component after nicotinic blockade with DHβE.

The isolated nicotinic opto-ACh current differed significantly and substantially with age-group (**Fig 2A,B**). The older group displayed ~30% of the younger nicotinic response in terms of both peak amplitude (unpaired *t*-test, *t*_27_ = 3.3, *P* = 0.003; **Fig 2C**), and charge transfer (unpaired *t*-test, *t*_27_ = 3.3, *P* = 0.003; **Fig 2D**). Under this condition, there was also a significant age-group difference in rising slope (unpaired *t*-test, *t*_27_ = 3, *P* = 0.005; **Fig 2E**). In a subset of recordings, the nicotinic identity of the residual optogenetic response was confirmed with application of selective antagonist DHβE which abolished the residual current response (paired *t*-test, *t_21_* = 4.8, *P* = 0.0001). The isolated nicotinic currents differ markedly by age-group.

**Figure 2.**
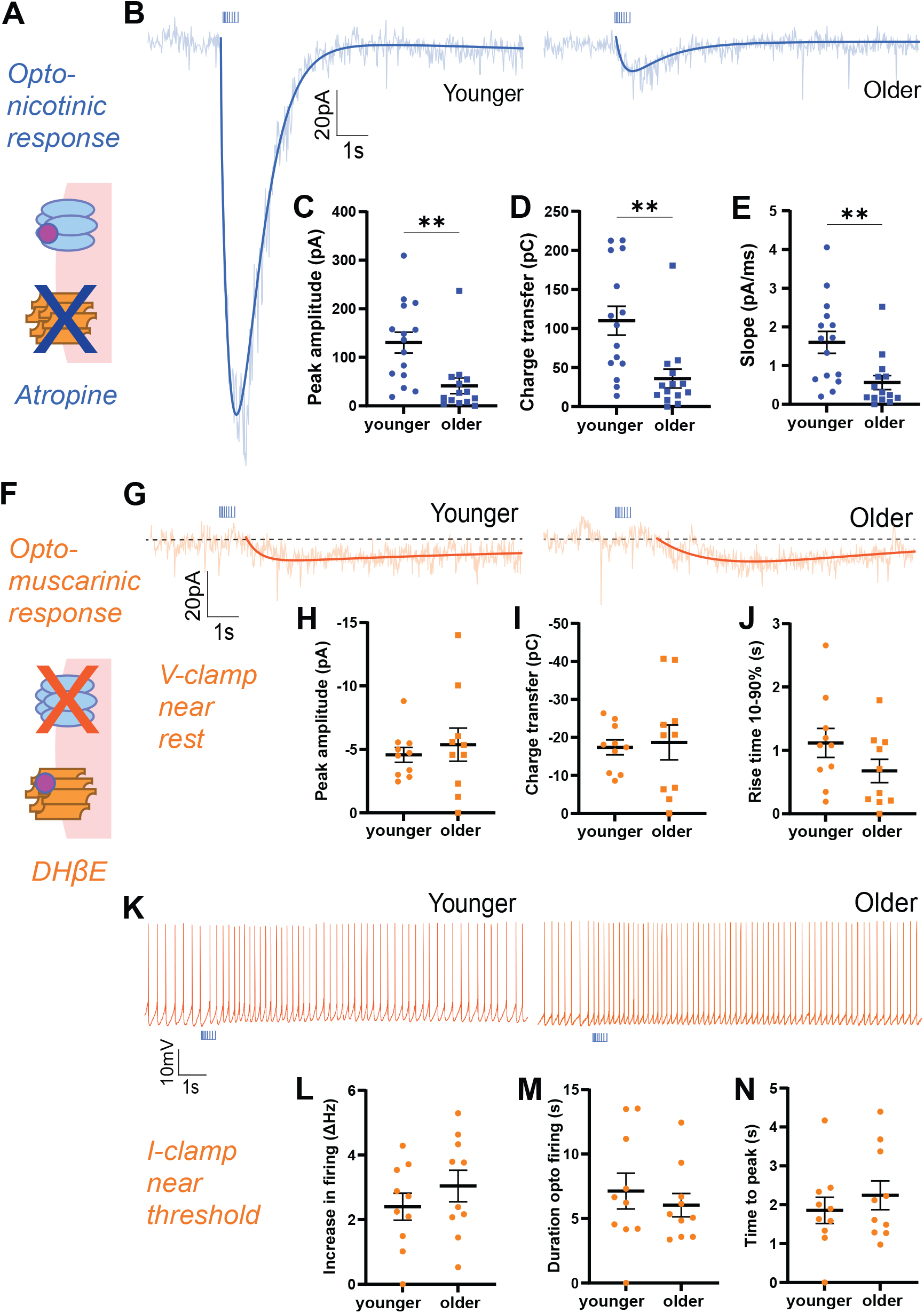
Nicotinic response is decreased and muscarinic response is preserved through adulthood. (**A**) Schematic illustrates muscarinic receptor blockade to isolate the nicotinic response. (**B**) Example optogenetically-elicited nicotinic responses recorded in voltage clamp (Vm = −75mV) in the presence of muscarinic receptor blocker atropine (AT) from younger and older mice. Graphs show significant age-group decreases in opto-ACh nicotinic responses: (**C**) peak amplitude, (**D**) charge transfer and (**E**) rising slope (\*\**P* <0.01, unpaired *t*-test, n = 29 neurons, 18 mice) (younger: P92 ± 5.2, P68-114; older: P184 ± 8.5, P150-226). (**F**) Schematic illustrates nicotinic receptor blockade to isolate the postsynaptic muscarinic response. (**G**) Example responses recorded in voltage clamp (Vm = −75mV) in the presence of nicotinic receptor blocker DHβE show muscarinic currents in neurons from younger and older mice. Graphs show no significant age group differences in opto-ACh muscarinic currents: (**H**) peak amplitude, (**I**) charge transfer, or (**J**) 10-90% rise time. (**K**) Example responses recorded in current clamp with depolarizing current injected to bring the neuron to fire action potentials in the presence of nicotinic receptor blocker DHβE show muscarinic firing response in neurons from younger and older mice. Graphs show no significant age-group difference in (**L**) peak firing frequency, (**M**) duration of opto-ACh firing change, or (**N**) time to peak firing (younger: P86 ± 20.2, P55-143; older: P236 ± 23.1, P167-303).

By contrast, the isolated muscarinic opto-ACh currents do not differ by age-group (**Fig 2F,G**). The older group displayed ~105% of the younger muscarinic response for peak amplitude (unpaired *t*-test, *t_18_* = 0.6, *P* = 0.6; **Fig 2H**) and ~95% for charge transfer (*t_18_* = 0.3, *P* = 0.8; **Fig 2I**). The kinetics of the slower muscarinic response was better captured by 10-90% rise time, which also did not show a significant age-group difference (*t_18_* = 1.5, *P* = 0.2; **Fig 2J**).

Since muscarinic receptors are state-dependent and exert stronger responses when the cell is depolarized, near threshold, or firing action potentials [42,43], we also investigated the impact of opto-ACh on postsynaptic neurons held in current clamp, with injection of current to elicit ~2-3 Hz action potential firing at baseline. Under these conditions, opto-ACh increased the frequency of action potential firing (**Fig 2K**), and this response was insensitive to application of nicotinic receptor blocker DHβE (peak: paired *t*-test, *t_18_* = 0.6, *P* = 0.6; duration: paired *t*-test, *t_18_* = 0.02, *P* = 0.9; data not shown). Opto-ACh firing response did not differ significantly between younger and older groups (peak: unpaired *t*-test, *t_18_* = 1, *P* = 0.3; **Fig 2L**; duration: *t_18_* = 0.7, *P* = 0.5; **Fig 2M**). There was also no significant age-group difference in time to peak firing response (*t_18_* = 0.8, *P* = 0.5, **Fig 2N**). These results in current-clamp are consistent with those in voltage-clamp, confirming that the optogenetic muscarinic response is preserved across age groups.

These results support the conclusion that a change in postsynaptic nicotinic receptors, not in muscarinic receptors, is predominantly responsible for the age-dependent decline in opto-ACh responses.

### Treatment impact of nicotinic positive allosteric modulation diminishes with age

Towards restoring a strong nicotinic signal, we apply a nicotinic positive allosteric modulator NS9283 [58,59] known to improve cognition in young animals [60,58]. The success of this treatment in older animals will depend on there being sufficient availability of postsynaptic nicotinic receptors for potentiation. We find that NS9283 (1μM) strongly potentiates younger opto-ACh responses, but this potentiation is less robust in older responses (**Fig 3A, B**) with a significant interaction of NS9283 and age on peak amplitude of opto-ACh (repeated measures two-way ANOVA, Interaction: F_1,24_ = 4.9, *P* = 0.04; **Fig 3C**). Sidak *post hoc* tests confirm significant NS9283 potentiation in the younger group (*t*_14_ = 5.1, *P* < 0.0001) that is not observed in the older group (*t*_13_ = 2, *P* = 0.1). Considering the NS9283 effect on opto-ACh charge transfer, there are significant main effects of age and drug (Two-way ANOVA, age: F1,24 = 9.2, *P* = 0.005; nicotinic PAM: F_1,24_ = 22.3, *P* < 0.0001; **Fig 3D**). Again, Sidak *post hoc* tests show NS9283 potentiation of opto-ACh charge transfer in the younger group (*t*_14_ = 4.7, *P* = 0.0001) and not the older group (*t*_13_ = 2, *P* = 0.1). This age-effect persists when comparing the percent increase with NS9283, where the increase in current amplitude in younger responses (57.1 ± 6.1%) is significantly larger than that seen in older responses (37.5 ± 8.4%; unpaired *t*-test: *t*_(24)_ = 2.2, *P* = 0.04). Improving nicotinic receptor function with NS9283 is effective in young adults but this efficacy diminishes in older adults. Our finding is consistent with behavioural work showing that NS9283 improves sustained attention, episodic memory, reference memory [60], and auditory discrimination [58] in younger animals. The decrease in effectiveness of the nicotinic PAM with age points toward a loss of nicotinic receptor availability. Therefore, we next investigate a candidate mechanism to improve the availability of nicotinic receptors.

**Figure 3.**
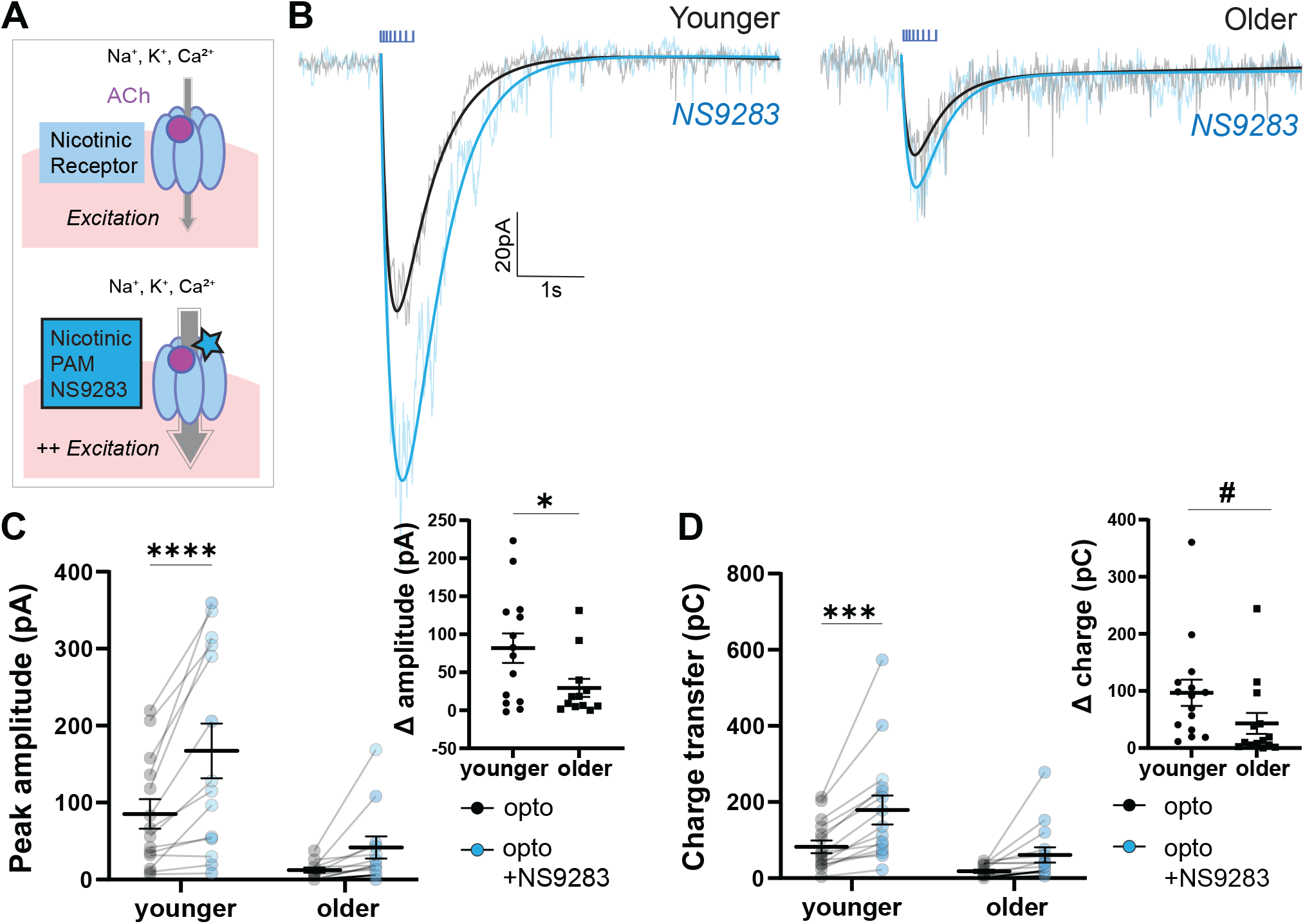
Older opto-ACh responses are less sensitive to nicotinic allosteric potentiation. (**A**) Schematics show acetylcholine activation of nicotinic receptors at baseline and with NS9283 positive allosteric modulation. (**B**) Paired recordings in voltage clamp (Vm = −75mV) show opto-ACh current responses before (black) and after (blue) bath-application of nicotinic positive allosteric modulator (PAM) NS9283 in neurons from younger and older mice. (**C**) Graph shows peak amplitude of paired opto-ACh currents before and after NS9283 in neurons from younger and older mice (\**P* < 0.05, Interaction, two-way ANOVA; \*\*\*\**P* < 0.0001, Sidak’s multiple comparisons). Inset graph shows the change in current amplitude imparted by NS9283 is significantly greater in younger than older mice (\**P* <0.05, unpaired t-test). (**D**) Graph shows charge transfer of paired opto-ACh currents before and after NS9283 in neurons from younger and older mice (\*\*\*\**P* < 0.0001, NS9283, \*\**P* < 0.01, Age, two-way ANOVA; \*\*\**P* < 0.001, Sidak’s multiple comparisons). Inset graph shows the change in charge transfer imparted by NS9283 trends towards a difference between younger and older mice (*t_24_* = 1.8, ^#^*P* = 0.08) (younger: P97 ± 12.1, P56-138; older: P227 ± 17.8, P166-295).

### Harnessing muscarinic receptor PKC signaling to improve nicotinic neurotransmission

The signaling pathways activated by muscarinic receptors [61–63] have independently been shown to improve nicotinic receptor insertion in the cell membrane [64,65]. To enhance nicotinic receptor availability, we therefore pursue an unorthodox strategy of stimulating muscarinic receptors to activate their secondary messenger pathways. We hypothesize that activating intact M1 muscarinic receptor signaling with direct agonist and cognitive enhancer xanomeline [66] will upregulate postsynaptic nicotinic receptor availability in older mice (**Fig 4A**).

**Figure 4.**
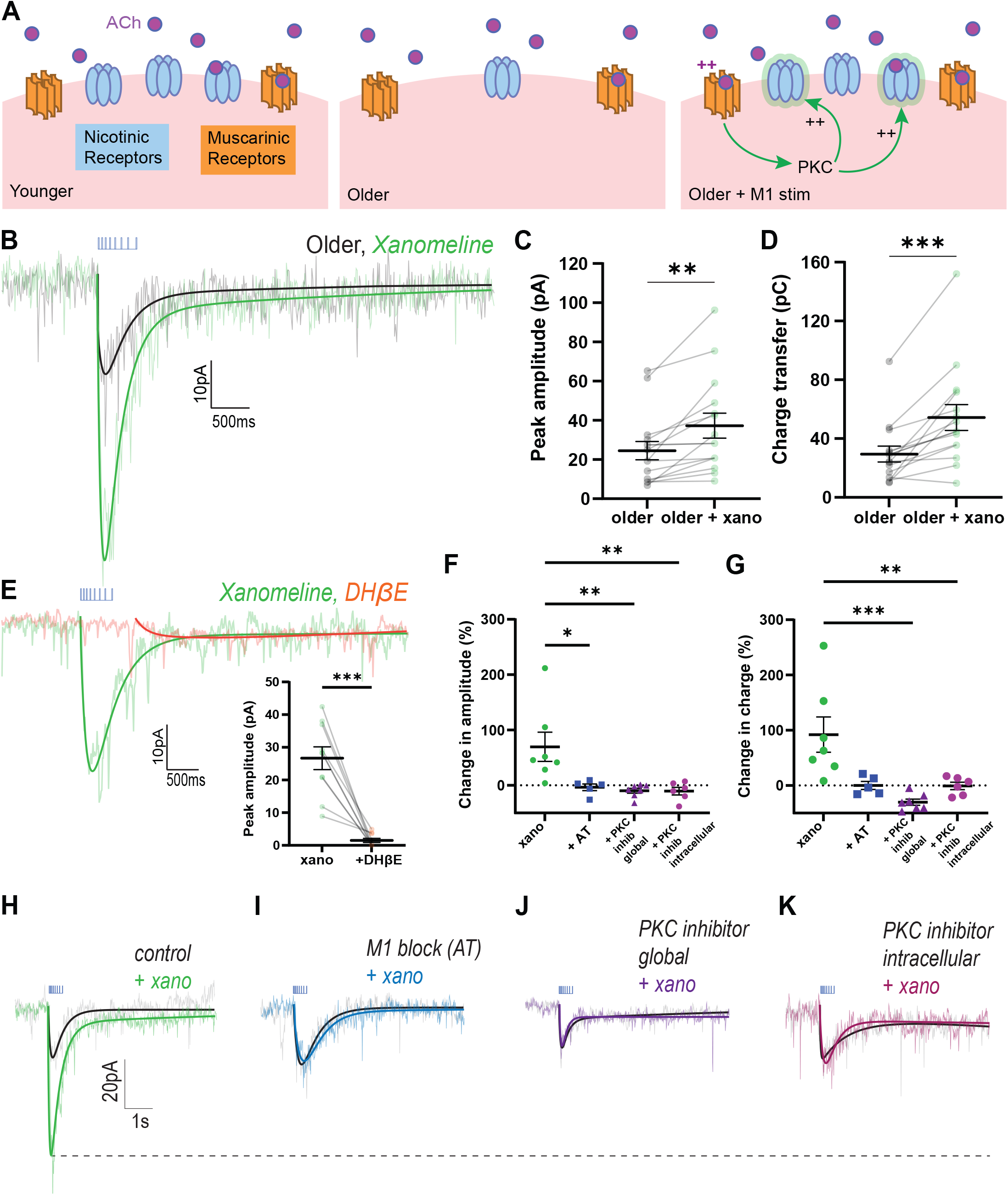
Older nicotinic responses improved by muscarinic M1 agonist xanomeline. (**A**) Schematic of cholinergic receptor activation in prefrontal layer 6 pyramidal neurons in younger, older, and older with intervention to improve nicotinic receptor availability. (**B**) Paired recordings in voltage clamp (Vm = −75mV) in older mice (mean: P223 ± 20.7 range: P150-358) show opto-ACh response before (black) and after (green) bath-application of muscarinic M1 agonist xanomeline. (**C, D**) Graphs show peak amplitude and charge transfer of older responses significantly increases with xanomeline (\*\**P* < 0.01, \*\*\**P* < 0.001, paired *t*-test). (**E**) Paired recordings in voltage clamp (Vm = −75mV) show opto-ACh response with xanomeline (green) is greatly reduced by nicotinic receptor blocker DHβE (orange). Graph (*inset*) shows peak amplitude of responses with xanomeline are significantly reduced with application of DHβE (\*\*\**P* < 0.001, paired *t*-test). (**F, G**) Graphs show potentiation with xanomeline is blocked in the presence of muscarinic receptor blocker atropine (AT), global PKC inhibitors (chelerythrine or Go6983), and intracellular PKC inhibitor (PKC 19-31). There is a significant drug effect on the percent change of peak amplitude, (\*\**P* < 0.01, one-way ANOVA; \**P* < 0.05, ** *P* < 0.01, Dunett’s *post hoc* tests) and charge transfer (\**P* < 0.05, one-way ANOVA; ** *P* < 0.01, Dunett’s *post hoc* tests). Example responses show xanomeline-potentiation of the opto-ACh response, (**H**) under control conditions, is blocked by pre-application of (**I**) muscarinic receptor antagonist atropine (AT), (**J**) global PKC inhibitor (Go6983), or (**K**) intracellular PKC inhibitor (PKC 19-31 peptide).

In prefrontal brain slices from older mice (P223 ± 20.7 days), application of xanomeline (300nM, 10 min) significantly increased opto-ACh responses (peak amplitude: paired *t*-test, *t_14_* = 3.2, *P* = 0.006; charge transfer: *t_14_* = 3, *P* = 0.01, (**Fig 4B**) with negligible impact on baseline holding current (−0.8 ±1pA). Following xanomeline treatment, we found opto-ACh responses continued to increase, with most neurons reaching a peak response 10-20 minutes post-xanomeline. Over time, xanomeline yielded a greater significant increase in opto-ACh peak amplitude (paired *t*-test: *t_14_* = 3.9, *P* = 0.002; **Fig 4C**) and charge transfer (paired two-tailed *t*-test: *t_14_* = 4.7, *P* = 0.0003; **Fig 4D**). For an example of this time course, see **Supplemental Figure S4**. In the absence of xanomeline, control experiments recorded over the same time course did not show potentiation of opto-ACh response (control change in peak with time: paired *t*-test, *t_14_* = 0.6, *P* = 0.6; control change in charge transfer with time: *t_14_* = 0.3, *P* = 0.8; data not shown). Consistent with our hypothesis that xanomeline boosts opto-ACh response by increasing the nicotinic component, nicotinic receptor antagonist DHβE eliminated the potentiated opto-ACh response (**Fig 4E**), significantly decreasing peak amplitude (paired *t*-test, *t_9_* = 6.6, *P* = 0.0001; **Fig 4E** *inset*) and charge transfer (*t_9_* = 4, *P* = 0.003, data not shown). These results demonstrate that xanomeline enhances the optogenetic nicotinic response in older mice.

To investigate the mechanisms underlying xanomeline-potentiation of the nicotinic component, we contrast the effects of xanomeline alone and after pathway manipulations in brain slices from the same mice (**Fig 4G,H**). We found that muscarinic receptor blockade with atropine eliminated xanomeline’s potentiation of opto-ACh responses, consistent with a requirement for activation of muscarinic receptors. Likewise, inhibition of PKC either globally, with bath-applied chelerythrine (1μM) or Go6983 (300nM), or only in the recorded neuron, with intracellular delivery of PKC 19-31 peptide (1μM) in the patch solution, also severely disrupted xanomeline potentiation of opto-ACh responses, consistent with dependence on PKC. Analyzing these experiments together, one-way ANOVA illustrates that all manipulations are sufficient to disrupt xanomeline potentiation of peak amplitude (F_3,21_ = 6.7, *P* = 0.002; Dunnett’s multiple comparisons, xano vs +AT: q_21_ = 3.1, *P* = 0.01; xano vs +PKC inhib global: q_21_ = 3.8, *P* = 0.003; xano vs +PKC inhib intracellular: q_21_ = 3.7, *P* =0.004 **Fig 4G,H**). **Figure 4** shows paired example responses illustrating that M1 inhibition, global PKC inhibition, and intracellular PKC inhibition prevent xanomeline-mediated potentiation (**Fig 4H-K**). Together, these experiments suggest that xanomeline-elicited potentiation requires downstream activation of intracellular PKC by muscarinic receptors, increasing the nicotinic component of the opto-ACh response in older adult mice.

### Full restoration with combined treatment of xanomeline and NS9283

To examine the sensitivity of the restored nicotinic response to further amplification, we applied xanomeline followed by NS9283 in brain slices from an additional group of older mice (P333 ± 6 days). To achieve a similar time-course to previous experiments with NS9283 (**Fig 3**), slices were treated with xanomeline then neurons were patched for paired recordings before and after application of NS9283 (**Fig 5A**). Application of nicotinic PAM after xanomeline yielded a highly significant increase in peak amplitude (paired *t*-test, *t_11_* = 5.3, *P* = 0.0003; **Fig 5B**), charge transfer (paired *t*-test, *t_11_* = 4.7, *P* = 0.0007; **Fig 5C**) and rise speed (paired *t*-test, *t_11_* = 5.7, *P* = 0.0001; **Fig 5D**). This striking potentiation of the older opto-ACh response by NS9283 further supports the conclusion that xanomeline increases postsynaptic nicotinic receptor availability.

**Figure 5.**
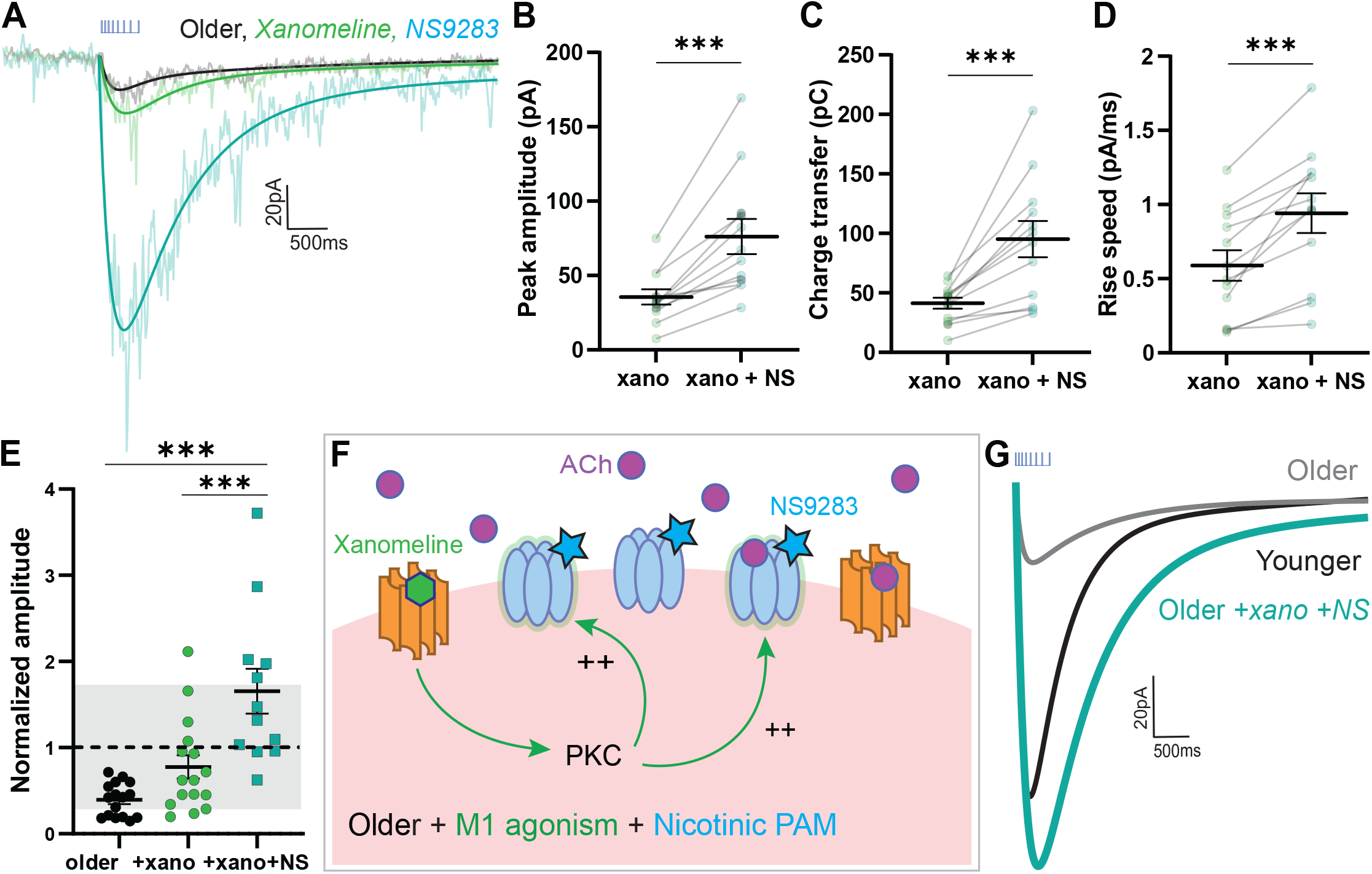
Older opto-ACh responses rescued to younger levels by combined treatment of M1 agonist xanomeline and nicotinic PAM NS9283. (**A**) Recordings in voltage clamp (Vm = −75mV) show the opto-ACh response in a neuron from an older mouse at baseline (black), with xanomeline (green), and with NS9283 (teal). In paired experiments, (**B**) Peak amplitude, (**C**) charge transfer, and (**D**) rising slope of older (mean: P333 ± 5.9, range: P325-344) xanomeline-enhanced responses (xano) significantly increase with NS9283 (\*\*\**P* < 0.001, paired t-test). (**E**) Graph shows peak amplitude of opto-ACh responses normalized to the mean peak amplitude of the younger opto-ACh responses, with grey denoting one standard deviation of younger opto-ACh responses. There is a significant effect of pharmacological interventions in rescuing older responses (\*\*\*\**P* < 0.0001, one-way ANOVA; *** *P* < 0.001, Tukey’s *post hoc* tests). (**F**) Schematic shows model for nicotinic receptor changes with age and the effects of combined xanomeline and NS9283 pharmacological intervention. (**G**) Example exponential fits of opto-ACh responses from younger, older, and older neuron treated with xanomeline and NS9283. Combined pharmacological intervention rescues older opto-ACh responses to younger levels.

Next, we normalized opto-ACh responses from this group of much older mice to the baseline mean of the younger mice (**Fig 5E**). This normalization illustrates the replication of the decline in opto-ACh response with age. It also demonstrates the ability of combination treatment of xanomeline with NS9283 to restore older responses into the typical range of young adults (one-way ANOVA, F_2,41_ = 16.49, *P* < 0.0001; Tukey’s *post hoc:* older vs xanomeline+NS9283: q_28_ = 8, *P* < 0.0001). Even in year-old mice, the deficit in opto-ACh signal can be restored to younger levels via combined treatment of xanomeline and NS9283 (**Fig 5F,G**). For further visualization of normalized responses with pharmacological intervention, see **Supplemental Figure S3**.

Overall, we present a novel strategy to restore nicotinic receptor neurophysiology in older mice informed by careful functional dissection of pre- and post-synaptic components of prefrontal cholinergic synapses.

## Discussion

Optogenetic activation of prefrontal cholinergic synapses reveals a significant decline in signaling during adulthood. Across the age range examined, multiple aspects of cholinergic neurotransmission are stable, with the critical exception of the decline in postsynaptic nicotinic receptor signal. While direct potentiation of nicotinic receptors with positive allosteric modulator NS9283 improves nicotinic responses in young adults, it imparts minimal benefit in older mice. We demonstrate that harnessing preserved muscarinic signaling via xanomeline upregulates nicotinic receptor availability in prefrontal cortex, restoring sensitivity to allosteric modulation. Combining xanomeline with nicotinic allosteric potentiation rescues the speed, strength, and receptor mechanism of endogenous cholinergic responses to younger levels.

### How age impacts pre- and post-synaptic aspects of the prefrontal cholinergic system

The net effect of age on cholinergic neurons and cortical signaling has been a subject of considerable debate. Multiple, sometimes conflicting, alterations in pre-and post-synaptic cholinergic components have been described during adulthood [40,67–74], leaving a knowledge gap in their integrated functional consequences. Here, we identify an age-related selective postsynaptic nicotinic receptor deficit in functional prefrontal cholinergic neurotransmission. This finding is consistent with earlier preclinical research showing age-dependent cortical decreases in nicotinic receptor binding [40,70,72,75] and in response to exogenous agonist [41]. A parallel decline in cortical nicotinic receptor binding is observed in older humans [17,18,76], underscoring the translational relevance of this work.

In contrast to the declining nicotinic signal, we found optogenetically-evoked postsynaptic muscarinic signaling remained stable across the age range studied. This stability was observed for two electrophysiological measures mediated by distinct cellular mechanisms [43,77–79], shedding new light on earlier research [67,80–83]. In contrast to prior reports at the whole tissue level [68,71,73,74], regulation of prefrontal cholinergic synapses by autoinhibition and acetylcholinesterase remain similar between age groups. These age-stable measures, together with successful rescue of nicotinic signaling, suggest presynaptic acetylcholine remains intact. In prefrontal cortex of healthy mice, there appears to be steady maintenance of trophic support mechanisms for cholinergic afferents over the age range we examined [84]. In healthy human aging, there are only gradual changes in cholinergic neurons and axons [85–87], with large or rapid declines in cholinergic afferents increasingly viewed as the domain of pathological aging [84,88,89].

### Implications for interventions to improve cholinergic signaling

Progressive decline in certain aspects of cognition during adulthood has been documented clinically [1–6] and preclinically [33–35]. Due to the importance of cholinergic signaling for executive function [7,8,10], acetylcholine receptors have been targeted extensively by therapeutic manipulations for cognitive improvement [23–29]. Our results indicate that improving nicotinic receptor signaling may allow for cognitive restoration. Direct stimulation of nicotinic receptors is complicated by desensitization [90–92]. Nicotinic receptor positive allosteric modulators, such as NS9283, present less risk of desensitization [93] and are pro-cognitive in young animals [58,60]. Our results (**Fig 3**) suggest the cognitive success of allosteric modulation in older subjects would be constrained and (**Fig 4**) highlight a strategy of first boosting nicotinic receptor availability via intracellular transduction pathways.

In cell systems, PKC upregulates nicotinic receptor trafficking [64,65] and phosphorylation [94]. This important kinase is activated by excitatory muscarinic receptors [61–63] and optophysiological experiments reveal proximity of muscarinic receptors to prefrontal cholinergic synapses (**Fig 2**) [36,39,79]. We reveal that activating muscarinic signalling via direct agonist xanomeline increases functional nicotinic responses in a PKC-dependent manner (**Fig 4**). This is consistent with a previous report that stimulating PKC through a different Gα_q_-coupled receptor increased electrically-evoked nicotinic responses without altering the probability of presynaptic acetylcholine release [95]. Here, we find xanomeline-upregulated nicotinic receptors restored youthful sensitivity of prefrontal cholinergic synapses to allosteric potentiation (**Fig 5**).

### Caveats and future research directions

Tools to examine cholinergic neurotransmission are not without limitations. Here, we discuss caveats for stimulus pattern and model selection. We employed optogenetic stimuli in a descending frequency pattern to mimic firing properties of the major subgroup of cortically-projecting cholinergic neurons [50–52]. Relatively little is known, however, about how these firing patterns change in adulthood, as previous work focused predominantly on younger animals [50–52]. There is evidence, however, that cholinergic firing patterns do not change substantially with age from juvenile to young adult [96]. Complications in cholinergic stimulation include limitations in achieving opsin expression in cholinergic neurons. The two major strategies in the field are suspected to alter cholinergic release in different directions (increased [97]; decreased [98]) and are associated with cognitive and behavioural changes from wildtype [97,98]. In previous electrophysiological work, we have examined the different models and found many similarities in their prefrontal cholinergic synapses in autoinhibition, metabolism, and postsynaptic receptor signaling [39], extending work from other groups [36,79]. While it remains possible that changes in ChAT promoter-driven opsin expression may interact with our findings, previous work has found that ChAT expression itself is well maintained over much of the age range used in this work [99], suggesting that changes in its promoter are unlikely to drive our reported age-effect. Here, our finding of age-related functional decline in nicotinic responses is consistent with work in wildtype rodents showing decreased nicotinic binding and diminished responses to exogenous agonist [40]. Furthermore, current discovery that xanomeline upregulates postsynaptic nicotinic signaling in a PKC dependent manner matches the consequences of electrical activation of a different cholinergic circuit in wildtype mice [95].

There are also caveats to consider in receptor and second messenger pathway specificity. Our results give new insight into a previously-unknown mechanism underlying xanomeline-mediated cognitive enhancement. The atropine-sensitivity and postsynaptic PKC-dependence of its nicotinic receptor upregulation points to an excitatory muscarinic mechanism, but xanomeline has also been shown to inhibit M4 receptors [100,101] and could potentially disinhibit glutamate release. Our data implicate the involvement of PKC, typically activated by Gαq-signaling, but this signaling also triggers activation of PLC [102] and may regulate GPI-anchored lynx family proteins [103] that also modulate nicotinic receptors [104,105] directly [106–108] or indirectly [107,109]. We find an improvement in apparent nicotinic receptor availability, however, the question of the density and localization of nicotinic receptors prior to-and after treatment remains outstanding. While the complex action of pharmacological intervention explored here warrants further investigation, our results demonstrate, for the first time, that muscarinic signaling can be harnessed to restore deficits in postsynaptic nicotinic receptors.

### Clinical relevance

Muscarinic agonism as a treatment for cognitive impairment has long been of interest. The direct M1 agonist xanomeline was first found to reduce cognitive impairments in older adults with Alzheimer’s disease in clinical trials 25 years ago, but presented adverse peripheral effects of muscarinic agonism [27,110]. Recently, xanomeline together with a peripheral blocker has been revisited in clinical trial, showing improved cognition in schizophrenia with less-severe adverse effects [111]. Another muscarinic agonist has also recently been shown to be pro-cognitive in pre-clinical work, tolerable in clinical trial, and improve human fMRI signals of hippocampal activation during spatial cue encoding [112]. These investigations highlight the importance of more fully understanding the mechanisms by which muscarinic signaling improves cognition.

### Summary of findings and implications

Our work investigates, for the first time, the functional consequences of aging on prefrontal cholinergic neurotransmission. We identify age-dependent functional deficits in postsynaptic nicotinic receptor activation. Using xanomeline to improve nicotinic signaling takes advantage of well-preserved muscarinic receptors and their second messenger pathways. Combining this treatment with nicotinic allosteric potentiation rescues endogenous nicotinic signaling. This reveals a novel mechanism for improving cognition through muscarinic receptors, taking advantage of a pharmacological agent already well investigated in clinical trials.

## Acknowledgments

We greatly appreciate the insight and feedback on earlier versions of this manuscript from Dr. Peter Carlen, Dr. JoAnne McLaurin, and Dr. Amy Ramsey of the University of Toronto.

## Author Contributions

Conceptualization, SKP, EKL; Methodology, SKP, SV, EKL; Data acquisition SKP, Analysis, SKP, SV, EKL; Writing, SKP EKL; Review & editing, SKP, SV, EKL; Funding, SKP, SV, EKL; Supervision, EKL. All authors have critically reviewed and approved of the manuscript.

