## Supplemental Methods, Table S1, Figures S1-S4 for "Xanomeline restores endogenous nicotinic acetylcholine receptor signaling in mouse prefrontal cortex"

### Supplemental Methods, Supplemental Table S1, & Supplemental Figures S1-S4

*Power et al.*

**Supplemental Methods**

*Animals*

Our colony was maintained by breeding on a C57BL/6 background. Male and female mice were weaned at postnatal day (P)21, separated by sex, and group housed (2-4 mice per cage) in plastic cages with corn cob bedding, houses for environmental enrichment, with *ad libitum* access to food and water on a 12hr light/dark cycle with lights on at 7:00AM.

*Electrophysiology*

Animals were anesthetized with an intraperitoneal injection of chloral hydrate (400mg/kg) and decapitated. The brain was rapidly removed in 4˚C sucrose ACSF (254mM sucrose, 10mM D-glucose, 26mM $NaHCO_3$, 2mM $CaCl_2$, 2mM $MgSO_4$, 3mM KCl, and 1.25mM $NaH_2PO_4$). The 400μm thick cortical slices of the prefrontal cortex (bregma 2.2-1.1) were obtained on a Dosaka linear slicer (SciMedia). Slices were transferred to a prechamber (Automate Scientific) where they recovered for at least 2hr in oxygenated (95% $O_2$, 5% $CO_2$) ACSF (128mM NaCl, 10mM D-glucose, 26mM $NaHCO_3$, 2mM $CaCl_2$, 2mM $MgSO_4$, 3mM KCl, and 1.25mM $NaH_2PO_4$) at 30˚C before being used for electrophysiology.

For whole-cell patch-clamp electrophysiology, brain slices were transferred to a chamber mounted on the stage of a BX51WI (Olympus) microscope and perfused with oxygenated ACSF at 30˚C at 3-4mL/min. Layer 6 pyramidal neurons were patched in accordance with their size, morphology, and proximity to white matter, as visualized using infrared differential interference contrast microscopy. Recording electrodes were filled with patch solution containing 120mM K-gluconate, 5mM $MgCl_2$, 4mM K-ATP, 0.4mM $Na_2$-GTP, 10mM $Na_2$-phosphocreatine, and 10mM HEPES buffer adjusted to pH 7.33 with KOH. Data were acquired and low-pass filtered at 20kHz with an Axopatch 200b amplifier (Molecular Devices) and Digidata 1440 digitizer and pClamp10.3 acquisition software (Molecular Devices).

*Pharmacology*

Atropine (200 nM; Sigma-Aldrich) was applied to block muscarinic receptors. Dihydro-β-erythroidine (DHβE; 3 μM; Tocris Bioscience) was applied to block β2-containing nicotinic receptors. CNQX (20 μM; Alomone Labs) and APV (50 μM; Alomone Labs) were used to block glutamate receptors. AF-DX 116 (300 nM; Tocris Bioscience) was used to block M2 muscarinic receptors. Galantamine hydrobromide (1 μM; Tocris Bioscience) was used to block acetylcholinesterase. Chelerythrine (10 μM; Tocris Bioscience) and Go6983 (300 nM; Tocris Bioscience) were used to block activation of PKC. PKC 19-31 peptide (1 μM; Millipore Sigma) was used to intracellularly block activation of PKC. It was aliquoted and frozen at 500 μM in 5% acetic acid and diluted to 1 μM in patch-solution on experiment day. NS9283(1 μM; Tocris Bioscience) was used to potentiate nicotinic responses. Xanomeline (300 nM; Tocris Bioscience) was used to activate M1 muscarinic receptors and increase nicotinic receptor availability. Unless otherwise stated, pharmacological agents were pre-applied for at least 10 minutes and co-applied during optogenetic stimuli and recovery period.

*Analysis*

Analysis was performed in Clampfit 10.3 (Molecular Devices) and Axograph. Raw traces were used for calculating rising slope of nicotinic current response within 50ms of opto-ACh onset to measure fast-onset kinetics. Downsampled traces were used to fit triple exponentials to cholinergic responses.

**Supplemental Table S1**

| | Capacitance | Input resistance | Resting membrane potential | Threshold | Spike amplitude |
|---|---|---|---|---|---|
| Younger < P150 (n = 109) | 86.6 ± 1.8 pF | 128.5 ± 3.2 MΩ | -87.3 ± 0.5 mV | 52.0 ± 0.8 mV | 67.7 ± 1.9 mV |
| Older ≥ P150 (n = 106) | 85.3 ± 2.0 pF | 125.7 ± 3.1 MΩ | -85.1 ± 0.6 mV | 50.5 ± 0.9 mV | 69.5 ± 1.7 mV |
| Statistics | $t_{(213)} = 0.5$, $P = 0.6$ | $t_{(213)} = 0.6$, $P = 0.6$ | **$t_{(213)} = 3$, $P = 0.004$ | $t_{(213)} = 1.2$, $P = 0.07$ | $t_{(213)} = 0.9$, $P = 0.4$ |

**Neuronal intrinsic properties by age group.** Table shows mean ± SEM for each intrinsic property in neurons from younger and older groups of mice, as well as the results of the unpaired *t*-tests.

**Supplemental Figure S1**

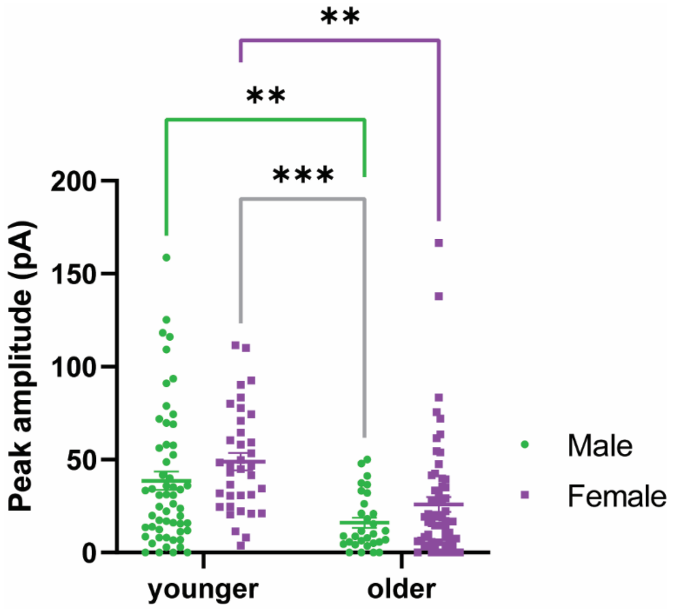


**Opto-ACh responses decrease with age in both sexes.** Graph shows opto-ACh response amplitude grouped by age and sex. There is no significant interaction between age and sex ($F_{1,180}$ = 0.002, $P$ = 0.97), but there is a highly significant age effect ($F_{1,180}$ = 23.5, $P$ < 0.0001) and a sex effect ($F_{1,180}$ = 4.6, $P$ = 0.03). Multiple comparisons analysis shows significant differences between younger and older responses for males (green circles, $t_{180}$ = 3.3, $P$ = 0.008) and for females (purple squares, $t_{180}$ = 3.6, $P$ = 0.003), as well as between younger females and older males ($t_{180}$ = 4.4, $P$ = 0.0001).

**Supplemental Figure S2**

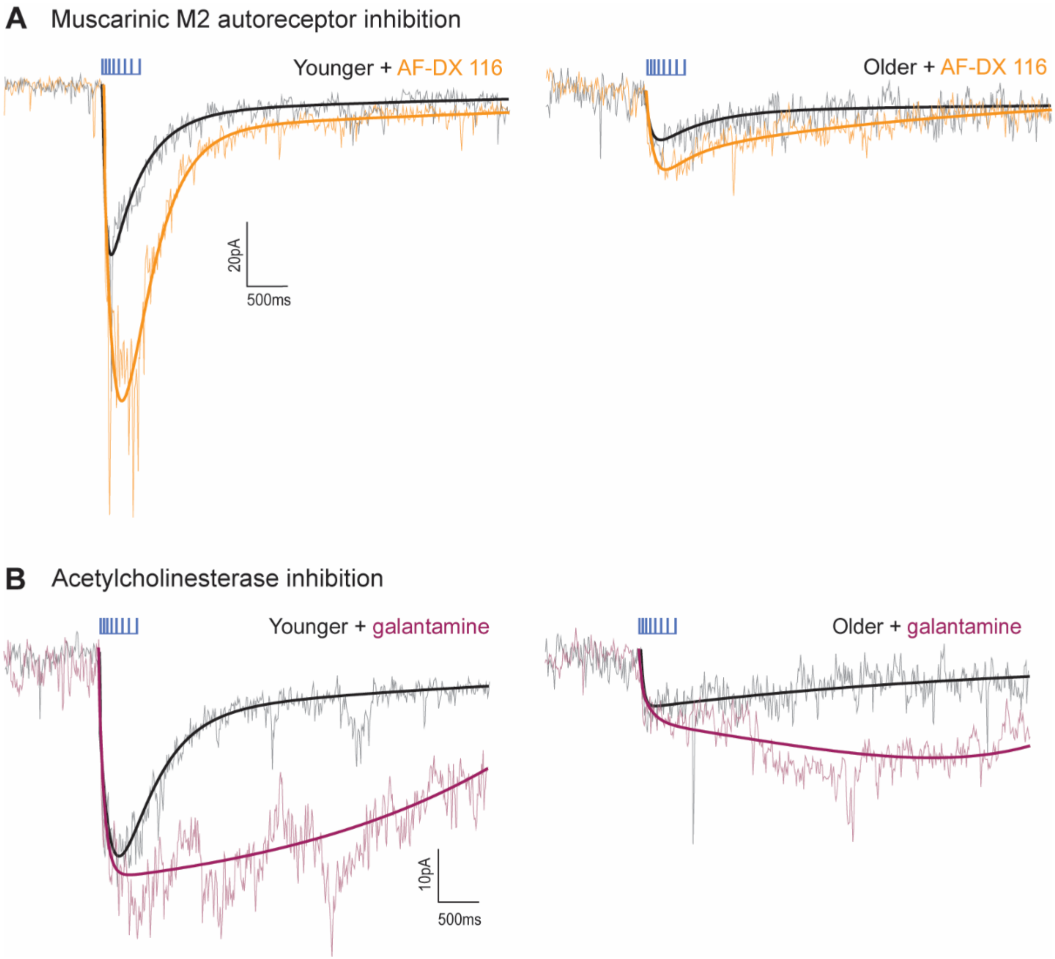


**Endogenous regulation of opto-ACh responses by autoinhibition and acetylcholine metabolism is similar across the age groups.** (**A**) Antagonist for M2 autoreceptor increases opto-ACh responses in both age groups: example paired responses before and after muscarinic M2 inhibition with AF-DX116 in a younger and older neuron. (**B**) Blocking acetylcholinesterase activity increases opto-ACh responses in both age groups: example paired responses before and after acetylcholinesterase inhibition with galantamine in a younger and older neuron. For quantitative investigation into these effects, see **Figure 1**.

**Supplemental Figure S3**

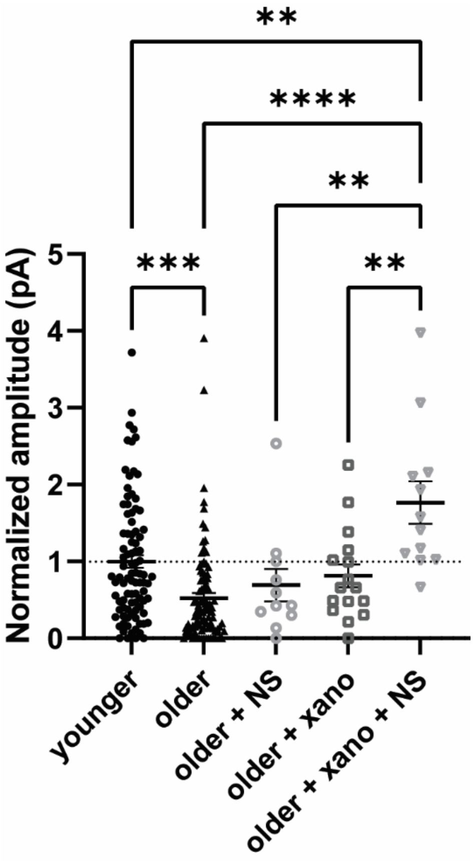


**Opto-ACh responses are increased by combination treatment of xanomeline + NS9283.** Graph shows opto-ACh response of older mice with amplitude normalized to mean response amplitude of the younger mice. Either NS9283 (NS) or xanomeline (xano) alone do not substantially improve older responses (changes are not significant as measured by multiple comparisons analysis), but combination of xanomeline + NS9283 substantially increases older responses, even beyond levels observed in younger mice (multiple comparisons $^{**}P < 0.01$, $^{***}P < 0.001$, $^{****}P < 0.0001$).

**Supplemental Figure S4**

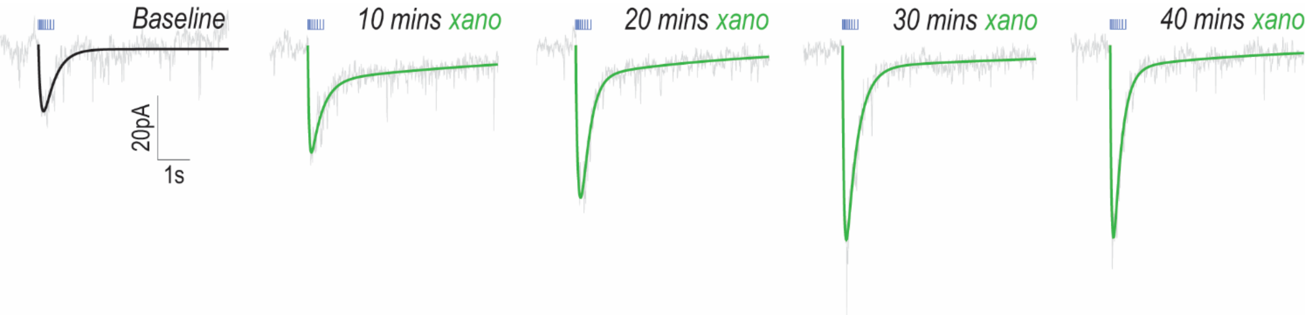


**Time course of xanomeline potentiation of opto-ACh response.** Paired response from one neuron showing time-course of xanomeline potentiation. Examples show baseline, immediately after bath-application of xanomeline for 10 minutes, and subsequent recordings every 10 minutes during washout, up to 40 minutes.
